# Reduced neural distinctiveness of speech representations in the middle-aged brain

**DOI:** 10.1101/2024.08.28.609778

**Authors:** Zhe-chen Guo, Jacie R. McHaney, Aravindakshan Parthasarathy, Bharath Chandrasekaran

**Affiliations:** Roxelyn and Richard Pepper Department of Communication Sciences and Disorders, Northwestern University, Evanston, IL, USA; Department of Communication Science and Disorders, University of Pittsburgh, Pittsburgh, PA, USA

**Author notes:** These authors contributed equally to this work (Zhe-chen Guo, Jacie R. McHaney).

**Keywords:** middle-aged adults, phoneme representation, neural dedifferentiation, cortical speech processing, electroencephalography

## Abstract

Speech perception declines independent of hearing thresholds in middle-age, and the neurobiological reasons are unclear. In line with the age-related neural dedifferentiation hypothesis, we predicted that middle-aged adults show less distinct cortical representations of phonemes and acoustic-phonetic features relative to younger adults. In addition to an extensive audiological, auditory electrophysiological, and speech perceptual test battery, we measured electroencephalographic responses time-locked to phoneme instances (phoneme-related potential; PRP) in naturalistic, continuous speech and trained neural network classifiers to predict phonemes from these responses. Consistent with age-related neural dedifferentiation, phoneme predictions were less accurate, more uncertain, and involved a broader network for middle-aged adults compared with younger adults. Representational similarity analysis revealed that the featural relationship between phonemes was less robust in middle-age. Electrophysiological and behavioral measures revealed signatures of cochlear neural degeneration (CND) and speech perceptual deficits in middle-aged adults relative to younger adults. Consistent with prior work in animal models, signatures of CND were associated with greater cortical dedifferentiation, explaining nearly a third of the variance in PRP prediction accuracy together with measures of acoustic neural processing. Notably, even after controlling for CND signatures and acoustic processing abilities, age-group differences in PRP prediction accuracy remained. Overall, our results reveal “fuzzier” phonemic representations, suggesting that age-related cortical neural dedifferentiation can occur even in middle-age and may underlie speech perceptual challenges, despite a normal audiogram.

## Introduction

Speech is a ubiquitous, socially relevant auditory signal that shapes spoken language communication throughout our lifespan. Despite its obvious value, speech perception can decline with age independently of audibility, a change that is distinctly noticeable by middle-age (1). Why do middle-aged adults show a decline in speech perception? Prior work has suggested that middle-aged adults may demonstrate temporal processing challenges due to peripheral cochlear neural degeneration (CND) (2), a condition that has been histologically verified in humans and animal models (3,4), but the perceptual consequences are still unclear and controversial (5). Here, we test a novel premise that middle-aged adults show reduced distinctiveness and greater dedifferentiation in the cortical representations of speech sounds that contribute to speech perceptual challenges. This premise is derived from the neural dedifferentiation hypothesis of aging, which posits that age-related processing challenges are driven by reduced specialization in the sensory cortex, leading to less distinctive and more correlated activity across the cortex (6–8). Ensembles in the temporal cortex encode critical phonological features that are causally linked to the percept of speech (9,10). Interestingly, these auditory cortical ensembles show age-related dedifferentiation and noisier processing due to induced CND in animal models (11,12). Given these findings, we asked whether cortical dedifferentiation of speech in middle-age relates to putative CND. In the next few sections, we expand on our understanding of cortical speech processing, the impact of aging on neural distinctiveness of speech, and the potential consequence of CND on cortical speech processing.

Speech perception has been variously described in the literature as being at the interface of neurobiology and linguistics and results from an interplay between bottom-up peripheral and top-down central processes (13). Yet, prior work examining speech perception in middle-aged adults (e.g., (2)) have almost exclusively focused on peripheral and subcortical neurobiology and peripheral-related declines, without alluding to the real possibility that some of the challenges could be driven by a disruption to cortical processes that are proximal to linguistically relevant operations. Patterns of cortical responses to simple speech syllables (e.g., /ba/ vs. /da/) are highly correlated with behavioral discriminability of these syllables in animal models, highlighting the proximity of cortical activity to speech perception (10). In real-world listening conditions, highly variable and continuous auditory signals need to be mapped to distinct, linguistically relevant, abstract phonological representations (14). Direct intracranial recordings using electrocorticography (ECoG) have revealed that phoneme representations are an emergent property of activity in distributed neuronal ensembles within the superior temporal gyrus (STG), which are selectively tuned to acoustic-phonetic features (9). For example, acoustic onset information is preferentially encoded in the posterior STG. In contrast, regions sensitive to phonetic cues are more spatially distributed along the posterior and anterior lateral STG (15,16) and show selectivity to phonetic features as granular as vowel height and consonant place (9). Phoneme representations emerge from the collective response patterns across these functionally distinct and anatomically interspersed local sites in the STG. Compared to younger adults, older adults exhibit less functionally specialized and distinct cortical representations (see (6) for a review). This decrease in cortical specialization is manifested as reduced category-selectivity of neural responses and has been noted in various domains, such as representations of visual stimuli

(8). In the domain of speech perception, an fMRI study by Du and colleagues (17) demonstrated age-related neural dedifferentiation of syllables, as evidenced by the finding that multivoxel pattern classifiers trained to classify phonemes performed less accurately for older adults than younger adults. These older adults also showed increased recruitment of frontal motor regions presumably to compensate for dedifferentiation of auditory representations. Currently, studies testing the neural dedifferentiation hypothesis of aging have focused on older adults, with little focus on middle-age (1). We hypothesize that dedifferentiation issues may already emerge in middle-aged adults without overt hearing loss, resulting in less differentiated cortical representations of phonemes.

Prior work has linked CND to auditory and speech discrimination deficits in animal models. By chemically inducing CND with ouabain, studies in mice provide direct evidence that CND increased the internal noise in the auditory cortex in a way that predicted the mice’s behavioral performance on selectively detecting target sounds in background noise (12). The increased cortical noise suggests that deficits in the auditory periphery can be one of the contributing factors of less dedifferentiated phoneme representations in human listeners. Therefore, in addition to examining age-related changes, it is crucial to examine the extent to which measures of putative CND—along with other peripheral, speech perceptual, and behavioral measures—may collectively explain variation in cortical phoneme differentiation.

To this end, we measured electroencephalographic (EEG) responses in younger and middle-aged adults with normal hearing thresholds while they listened to continuous speech in quiet (Fig. 1A). This paradigm probed the functionally and anatomically distinct response patterns in the auditory cortex evoked by continuous speech (18–20). We computed and analyzed “phoneme-related potentials” (PRPs) by averaging EEG responses time-locked to phoneme instances (Fig. 1C) to capture phoneme-specific neural dynamics organized by phonetic features, such as manner of articulation (21). We also measured neural tracking of basic, continuous acoustic properties, including acoustic envelope and onsets (Fig. 1B), which can change as a function of age (22,23) and drive differences in phoneme encoding. In addition to traditional audiometry, we also examined extended high frequencies, as elevated extended high-frequency hearing thresholds can impact speech perception (24), and auditory brainstem responses (ABR) as an objective physiological measure of sound processing along the auditory pathway (25). Specifically, we examined the amplitude of wave I as a proxy of the health of Type-I spiral ganglion neurons, as wave I amplitude is directly proportional to the quantity of intact synapses (4,26,27). Additionally, we also examined wave V as a proxy for homeostatic gain normalization, which has been previously evidenced as a marker of CND (28). Finally, we quantified speech perception performance using the Words in Noise test, which provides less linguistic scaffolding relative to other clinical tests (29), and two measures of cognition to rule out any impacts of cognition on speech perception abilities (30).

**Figure 1.**
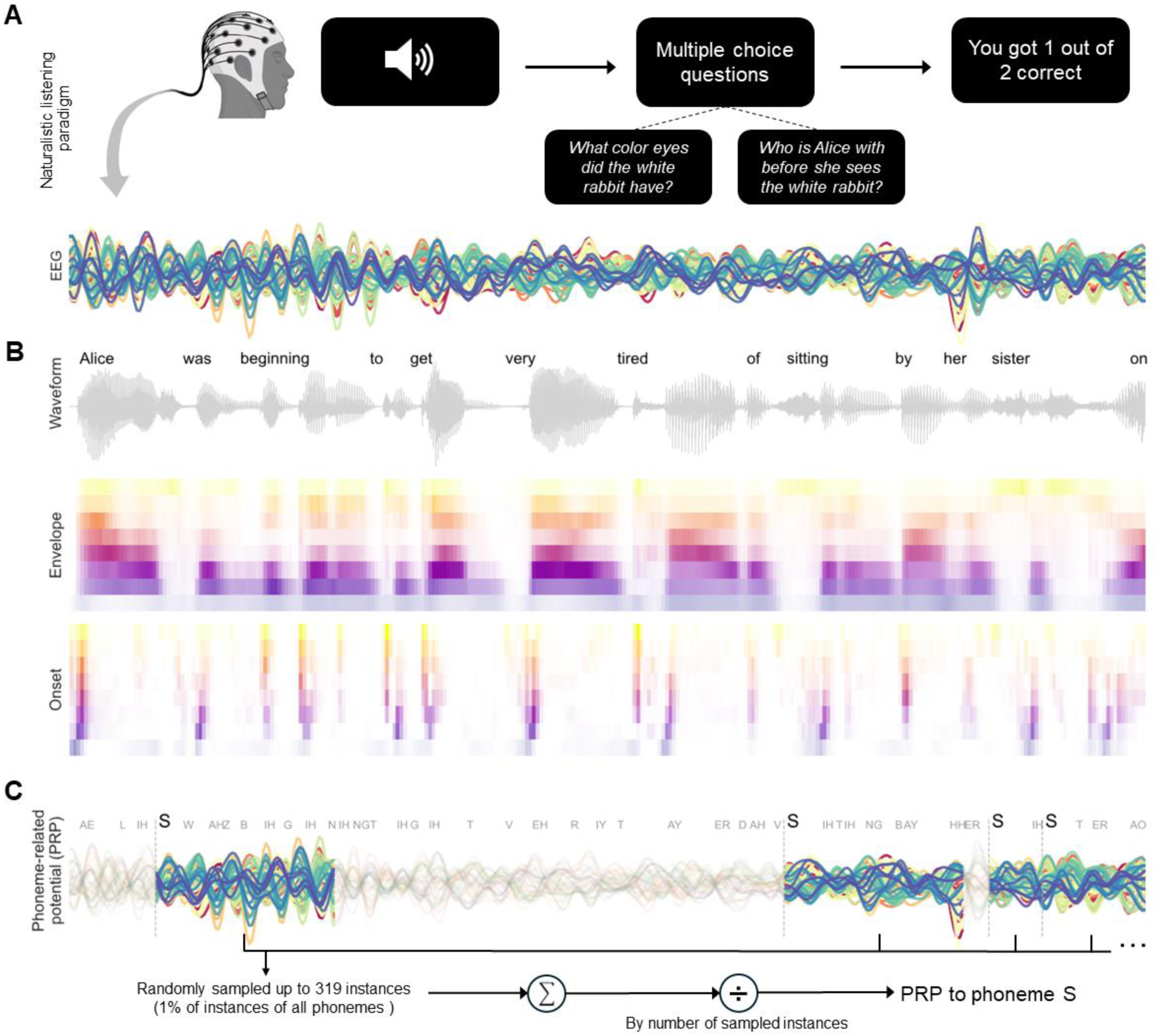
Continuous speech listening paradigm. A) Participants listened to the audiobook version of Alice’s Adventures in Wonderland while EEG activity was recorded. The figure shows preprocessed EEG responses from one participant time-aligned to an audiobook excerpt. At the end of each story segment (∼60 s), participants answered two comprehension questions regarding the content of the segment. B) Speech waveform and 8-band envelope and onset representations of the stimuli. C) For each participant, phoneme-related potentials (PRPs) were calculated as the averaged EEG responses from the onset of each phoneme instance (e.g., /s/) to 500 ms post-onset.

To anticipate, our findings demonstrate that cortical representations of phonemes and their featural relationship do become less distinct in middle-age, highlighting the role of cortical aging in speech perception. Signatures of putative CND were strongly associated with phoneme dedifferentiation. Yet, after controlling for CND signatures and other predictors, such as acoustic processing abilities, middle-aged adults still exhibited less distinct phoneme representations.

## Results

### Middle-aged adults showed less neural distinctiveness of phoneme encoding than younger adults

Participants in this study were recruited from a larger, cross-species study examining auditory and speech processing in middle-age that required five sessions of participation. From the larger participant pool, 44 participants completed the continuous speech listening task (Fig. 1). Of the 44 participants, 24 were younger adults (18–25 years, *M* = 21.4, *SD* = 2.02; 22 females) and 20 were middle-aged adults (40–54 years, *M* = 46.05, *SD* = 4.4; 12 females). All participants were required to have clinically normal pure tone air conduction thresholds, as defined by thresholds less than or equal to 25 dB HL at octave frequencies 0.25, 0.5, 1, 2, and 4 kHz.

We examined the extent to which younger and middle-aged adults differed in the cortical representations of phonemes by using PRPs derived from participants’ EEG responses to continuous speech in quiet. First, we analyzed neural separability of phonemes by computing the *F*-statistic on the PRPs over time (21) (Fig. 2A). Here, the *F*-statistic was the ratio of between-phoneme variability over within-phoneme variability, with higher values indicating greater category separation. Throughout the duration of the PRP, the *F*-statistic was generally higher for younger adults (*M* = 1.540, average *SD* = 0.472) than for middle-aged adults (*M* = 1.314, average *SD* = 0.316), especially during the first 350 ms, providing initial evidence for reduced phoneme separability in the middle-aged group.

**Figure 2.**
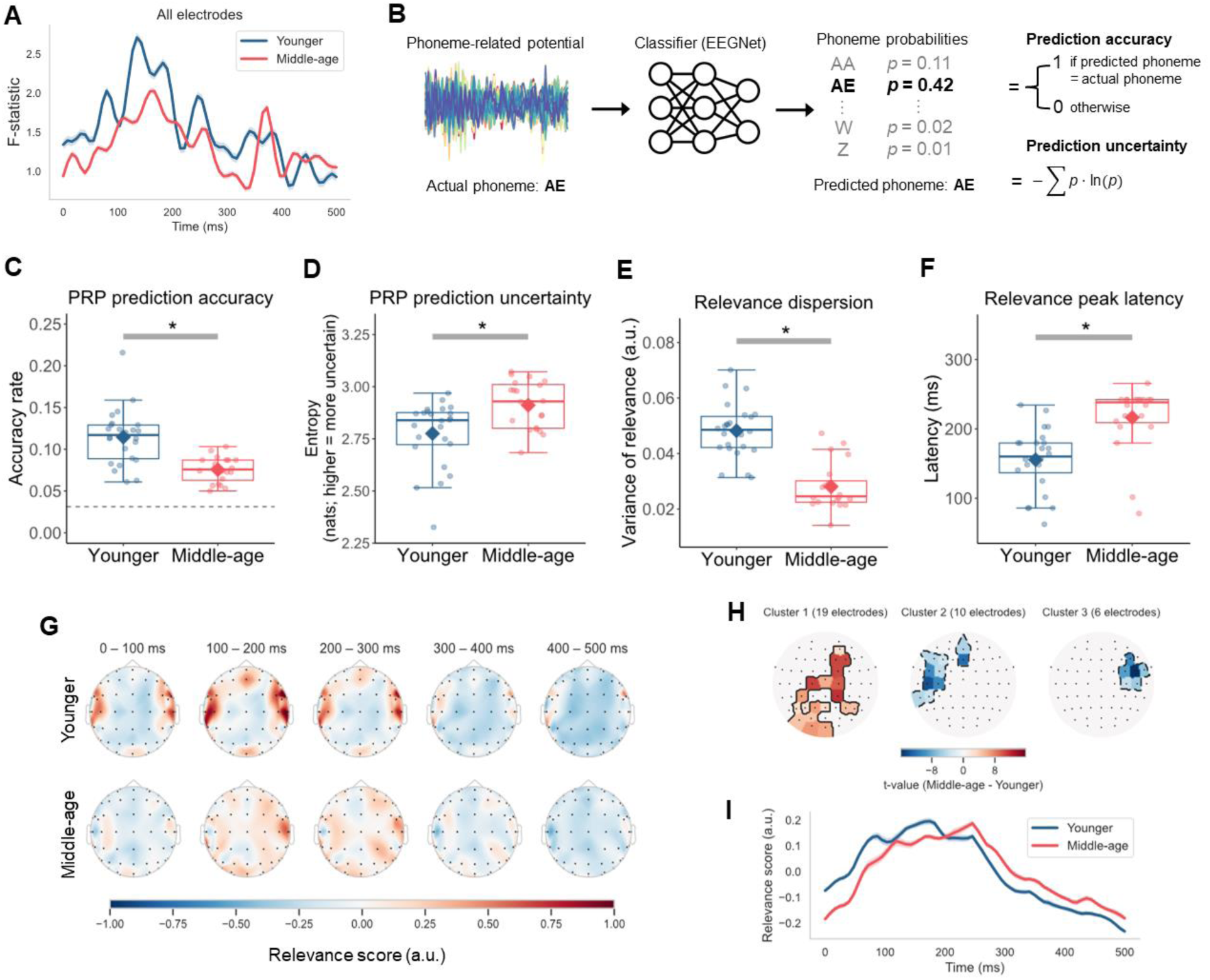
Less distinct phoneme encoding in middle-aged listeners. A) F-statistic of the PRPs over time calculated using all electrodes. Ribbons around the curve represent ±1 standard error. B) EEGNet convolutional neural network classifiers, trained with 20-fold cross-validation, predicted the phoneme label for each held-out PRP. The phoneme with the highest probability was recorded as the prediction and Shannon’s entropy measured the prediction uncertainty (higher entropy indicates more uncertainty). C) to F) Boxplots of phoneme prediction accuracy, prediction uncertainty, relevance dispersion, and relevance peak latency for the younger and middle-aged adults, with individual participants’ values (dots) and the group mean (diamond). Boxplots show the median (horizontal line), 25th and 75th percentiles (box edges), and ±1.5 interquartile range (whiskers). Asterisks indicate statistical significance at the 0.05 level. Dashed horizontal line in the accuracy panel marks the chance-level performance. G) Topographies of averaged relevance score in 100-ms time bins over the duration of PRP. H) Clusters of electrodes with significant group difference (p < 0.05, two-tailed) based on mass-univariate independent-samples t-tests comparing relevance scores (averaged over time) between younger and middle-aged adults. Electrodes are color-coded for t-values. I) Time course of relevance averaged over all electrodes and participants in each age group. Ribbons around the curve represent ±1 standard error.

To confirm that phoneme categories were less separable for the middle-aged group, we decoded the multidimensional PRP time series using the EEGNet classifier (31), a compact convolutional neural network specifically designed for classifying EEG signals across different brain-computer interface paradigms (Fig. 2B). A key advantage of neural networks like EEGNet is the ability to capture potentially non-linear dynamics that may integrate information from multiple distributed electrodes and demonstrate phoneme representations. Cross-validated EEGNet models were trained to predict phoneme labels for out-of-sample PRPs. We measured prediction accuracy and uncertainty, evaluated as Shannon’s entropy with higher values indicating more uncertain predictions.

The median phoneme prediction accuracy for both age groups (Fig. 2C) was significantly above the chance level of 3.23% (younger: *z* = 4.286, *p* < 0.001; middle-aged: *z* = 3.925, *p* < 0.001). Importantly, phoneme prediction accuracy of middle-aged adults (*Mdn* = 0.075) was significantly lower than that of younger adults (*Mdn* = 0.117; *z* = −4.151, *p* < 0.001), in line with the overall pattern of the *F*-statistic curves (Fig. 2A). Consistently, the phoneme prediction uncertainty of the PRP classifier was significantly greater for middle-aged adults (*Mdn* = 2.930) than for younger adults (*Mdn* = 2.839, *z* = 2.569, *p* = 0.010; Fig. 2D). These findings converge to suggest less distinct representations of phonemes in middle-age.

### Middle-aged adults’ phoneme processing involved a more distributed network with a delayed timing

We next investigated the extent to which spatio-temporal features of the PRP were relevant to phoneme identity and whether the patterns differed between the two age groups. We leveraged the ability to interpret the learned representations from EEGNet using the Deep Learning Important FeaTures (DeepLIFT) algorithm (32). DeepLIFT assigned a contribution score quantifying how much evidence each feature in the PRP provided towards the prediction of the actual phoneme class. We used the DeepLIFT contribution scores to derive relevance scores summarizing the relative degree to which each electrode contained information relevant to the target phoneme over time for each participant, with higher values representing greater relevance.

Topographic maps of average relevance scores in 100-ms intervals over the duration of the PRP for each age group are depicted in Figure 2G. One immediate observation was that the relevance was more localized for younger adults but more uniformly distributed for middle-aged adults. To test this observation, we averaged relevance scores over time for each electrode and computed relevance dispersion, or the variance of these scores between the electrodes, for each listener. Relative to that of the younger group (*M* = 0.048, *SD* = 0.010), the relevance dispersion of the middle-aged group was significantly smaller (*M* = 0.028, *SD* = 0.009; Welch’s *t*(41.851) = −7.213, *p* < 0.001, 95% CI[−0.026, −0.014]), suggesting that relevance values across electrodes were more similar, and hence, that important information for phoneme identity was more spatially distributed (Fig. 2E). We also conducted a mass-univariate independent *t*-test to specifically identify which electrodes showed significant group differences. This was a cluster-based permutation test that used uncorrected *p* ≤ 0.05 as the cluster forming threshold. Then, corrected *p*-values were computed for each cluster based on the cluster-mass statistician null distribution from 10,000 permutations (33). The results revealed three clusters of electrodes (Fig. 2H), including one large cluster spanning electrodes from the occipital to fronto-central area with higher relevance scores for middle-aged adults (*t_max_* = 11.573, *p* < 0.001). The other two smaller clusters included mostly temporal electrodes in the left (*t_max_* = −13.947, *p* < 0.001) and right (*t_max_* = −15.854, *p* < 0.001) hemisphere with lower relevance for the middle-aged group.

In addition to spatial dispersion, we compared the temporal dynamics of relevance scores across both age groups. For each participant, we averaged relevance over all electrodes at each time point in the PRP to derive a time-varying relevance curve (Fig. 2I) and identified the latency at which the relevance value reached its maximum. The relevance peak latencies of middle-aged adults (*Mdn* = 238.281) were delayed relative to those of younger adults (*Mdn* = 160.156; *z* = 4.208, *p* < 0.001; Fig. 2F). These results indicated that phoneme processing in middle-age may be less efficient and supported by a relatively broader cortical network.

### Reduced encoding of featural relationship between phonemes in middle-age

The above analysis treated phonemes as independent, atomic categories. Yet, cortical phoneme representations are an emergent property of neural ensembles selectively tuned to specific acoustic-phonetic features (9,15,16), which provide critical cues to phonemes, and hence, are fundamental to speech perception. It is possible that the reduced phoneme encoding in middle-age is also reflected in the feature-level connection between phonemes, which can be formally expressed using binary-valued distinctive features from phonological theories (34). In phonology, a phoneme can be specified with either + or – for a feature, indicating the presence or absence of that feature. The feature values collectively define the identity of a phoneme and classes of phonemes sharing similar properties. Given that cortical oscillations captured by EEG primarily reflect distinctions based on manner of articulation (21), we considered the three features that capture major manner class distinctions, such as vowels versus consonants: [syllabic], [sonorant], and [continuant] (Fig. 3A). [syllabic] indicates whether a sound can function as the syllable nucleus and distinguishes vowels ([+syllabic]) from consonants ([-syllabic]). [sonorant] refers to sounds produced with an open vocal tract and continuous non-turbulent airflow, distinguishing obstruents ([-sonorant]) versus approximants and nasals ([+sonorant]) among consonants. [continuant] specifies whether a sound is produced with the blocking of the oral cavity and differentiates between approximants ([+continuant]) versus nasals ([-continuant]) among sonorants and between fricatives ([+continuant]) versus stops ([-continuant]) among obstruents. Note that as instances of the same phoneme generally share the same values for the three manner-defining features regardless of their position in a word or variations in acoustic realization, the features model an abstract relationship between phonemes. We investigated the extent to which the participants’ PRPs reflect such a featural relationship through the representational similarity analysis (35).

**Figure 3.**
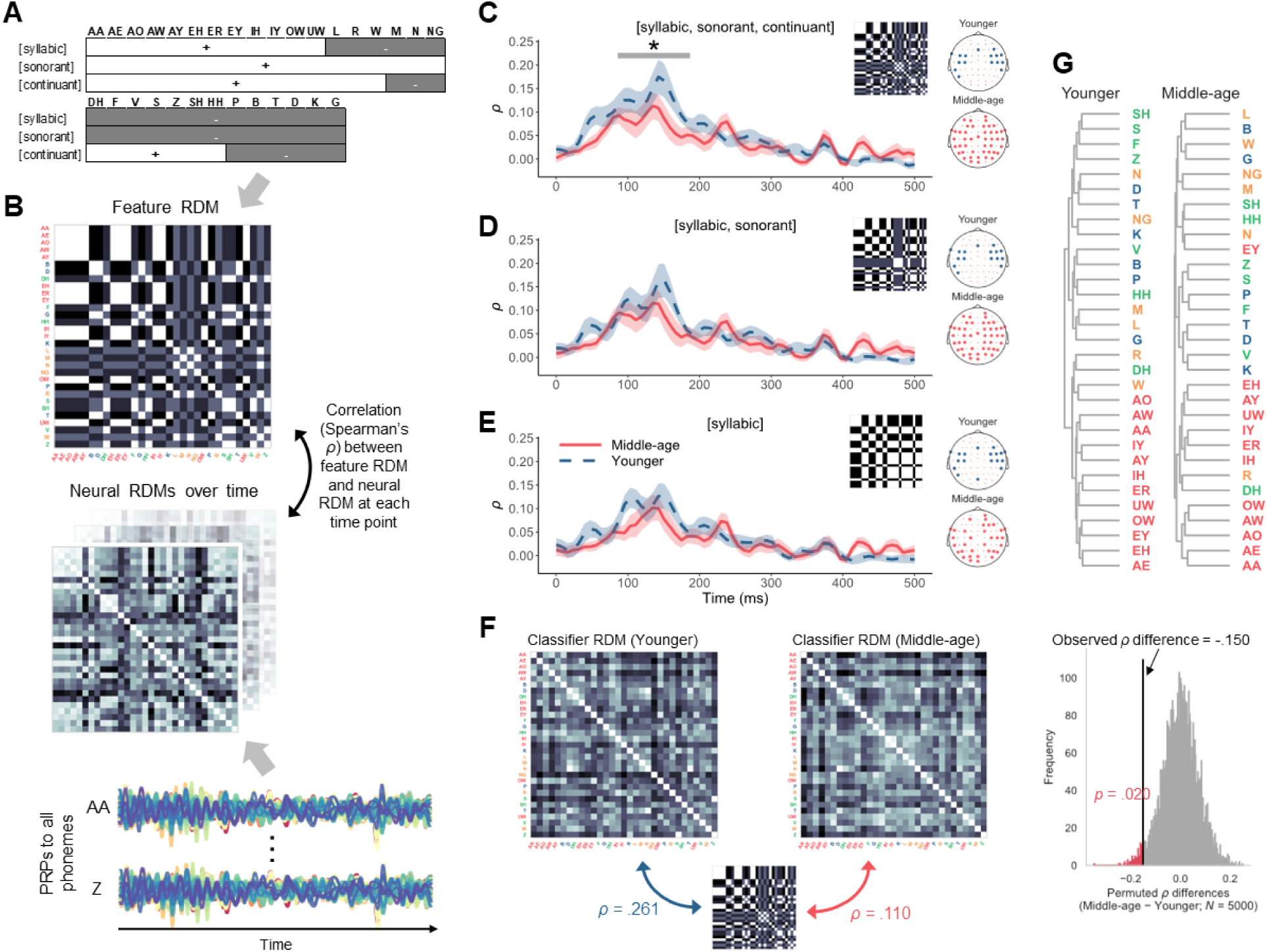
Less distinct featural relationship between phonemes in middle-age. A) Feature specification of the phonemes for [syllabic], [sonorant], and [continuant]. B) Phoneme dissimilarities based on the featural characterization in panel A were modeled as a representational dissimilarity matrix (RDM), which contains pairwise dissimilarity values (Euclidean distances) of the phonemes. Phoneme labels are colored based on the manner of articulation (vowels, approximants and nasals, fricative, stops). From the PRPs, we constructed a series of RDMs representing neural dissimilarity between phoneme pairs over time. The neural RDM at each time point was then compared with a feature RDM using Spearman’s rank correlation coefficient (ρ). C to E) Average correlation ρ of each age group over time with ribbons indicating ±1 standard error. Each panel presents the results for a different level of feature distinction. The topographies on the right show the electrodes included to build the neural RDMs, which were those leading to optimal alignment (highest overall ρ) with the RDM of each feature hypothesis. F) RDMs were derived from the confusion matrices of the PRP classifiers and compared with the feature RDM including [syllabic, sonorant, continuant] for each age group. The ρ difference between the younger and middle-aged groups was significant according to a permutation analysis which calculated a permuted p-value from a null distribution of ρ differences. G) Hierarchical clustering of the phonemes for each age group using PRPs during the 0– 350 ms time window and electrodes in panel C.

We constructed a representational dissimilarity matrix (RDM) that contained pairwise dissimilarity values for all 31 phonemes based on their feature values for [syllabic], [sonorant], and [continuant] (Fig. 3B). Similarly, from the PRPs, we derived a neural RDM at each time point for each participant. The neural RDMs over time were then compared against the feature RDM using Spearman’s correlation *ρ* as the similarity measure. Higher *ρ* indicated stronger alignment between neural and featural representations. In creating the neural RDMs for each age group, we sorted the electrodes of the PRPs by the relevance scores from the PRP classifiers and included the set of electrodes that maximized the correlation with the feature RDM.

For both age groups, the neural RDMs showed a correlation peak at approximately 150 ms after phoneme onset (Fig. 3C), similar to the timing of the phoneme separability (*F*-statistic) peak (Fig. 2A). Crucially, generalized additive modeling revealed that the correlation curve was significantly lower for middle-aged adults between 86–187 ms, suggesting poorer agreement with the feature RDM. We additionally tested two reduced feature RDMs with [syllabic] and [sonorant] and with only [syllabic] (Figs. 3D and 3E). There was no significant age-group difference when using these two reduced feature models. The findings suggested that cortical representations of phonemes are less distinct at the featural level in middle-aged listeners. Additionally, it is worth noting that the number of electrodes included to build the neural RDMs (topomaps on the right in Figs. 3C through 3E), which formed the set leading to the highest *ρ*, was greater for middle-aged adults than for younger adults, regardless of the feature RDM. Such a pattern indicated that information from a broader network of electrodes was required to achieve optimal alignment with the feature-based phoneme representations, consistent with the more distributed relevance from the previous analysis (Fig. 2G).

Finally, we found that the group difference in alignment with the featural representations of the phonemes was reflected in the confusion patterns of the PRP classifiers. The confusion matrices of the PRP classification from each group were used to construct RDMs, which were again compared with the full feature RDM containing [syllabic], [sonorant], and [continuant] through Spearman’s *ρ* (Fig. 3F). The correlation was higher for younger adults (*ρ* = 0.261) than for middle-aged adults (*ρ* = 0.110), and this difference was significant, according to a null distribution of *ρ* differences derived through a permutation analysis (Fig. 3H). Thus, not only was the encoding of individual phonemes less distinct in middle-age, but also their featural relationship was less preserved.

The results thus far are in line with the hypothesized cortical dedifferentiation of phonemes in middle-age. Meanwhile, we also assumed that the dedifferentiation may stem at least partially from 1) reduced cortical tracking of critical acoustic features associated with speech intelligibility, and 2) putative markers of CND. In the next sections, we examined these possibilities. Finally, to obtain a more holistic view of the age-related changes and the interplay between peripheral and central processes, we examined the extent to which these variables explain the neural differentiation of phoneme information.

### No evidence for age-group difference in the neural tracking of acoustic envelope and onsets

We first considered a more parsimonious, acoustic-based explanation for the age-group differences observed above—that middle-aged listeners were simply less capable of tracking continuous acoustic properties that are relevant for speech perception. It is well-documented that responses in the auditory cortex robustly track speech envelopes (36,37) and are particularly sensitive to acoustic edges (38,39). The ability to track these acoustic features is impacted by aging, with some evidence for a decline in this ability in older adults compared with younger adults (22), while other studies reported an increase (23). Here, we compared the cortical tracking of acoustic spectrogram (i.e., envelopes) at different frequency bands and the corresponding acoustic onset representations across the younger and middle-aged groups (Fig. 1B). The envelope and onset representations of the stimuli were included as the predictors in linear encoding models that predicted held-out EEG responses at each electrode through multivariate temporal response function (40) (mTRF) (Fig. 4A). Pearson’s correlations (*r*) between predicted and actual EEG responses served as the measure of neural tracking prediction accuracy. To assess changes in neural tracking strength that could be uniquely attributed to either the envelope or onset predictors alone, we subtracted the prediction accuracy of a model without the predictor of interest from that of a full model including both predictors.

**Figure 4.**
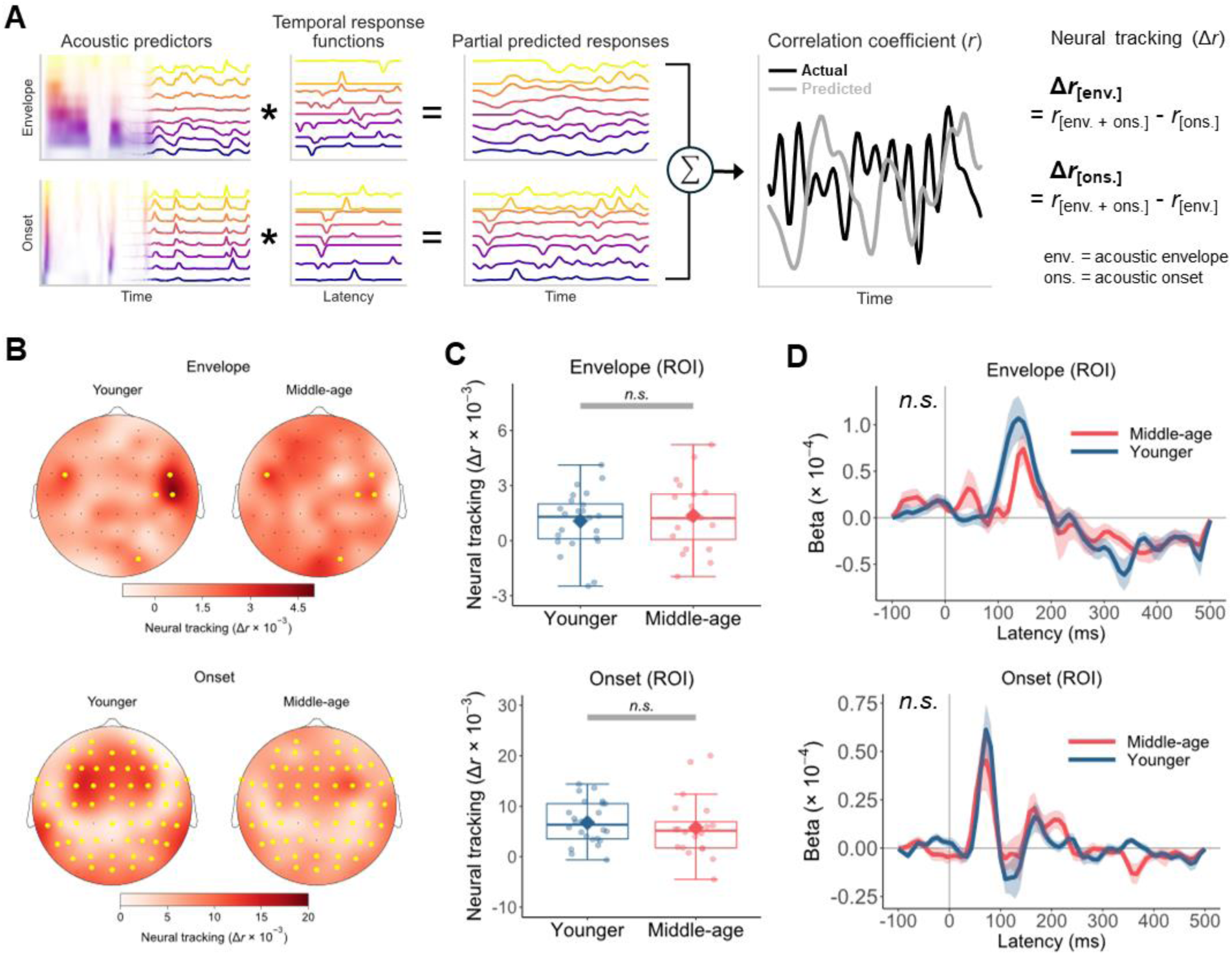
Estimation of neural tracking of acoustic envelope (spectrogram) and onset features. A) Amplitude envelope and onset envelope at each frequency band were convolved the corresponding cross-validated temporal response functions (TRFs) to predict partial neural responses, which were summed to derive the predicted response. The prediction was correlated with the actual EEG at each electrode using Pearson’s r as the accuracy metric. To isolate the neural tracking that could be attributed uniquely to either the envelope or onset predictor (Δr), we subtracted the prediction accuracy of the model without either predictor from that of the full model including both predictors. B) Topography of average neural tracking of envelope and onset features in younger and middle-aged listeners. The region of interest (ROI; yellow dots) was defined as the set of electrodes with neural tracking significantly above zero (p < 0.05, one-tailed) based on mass-univariate one-sample t-tests including all participants and collapsing across age groups. C) Individual participants’ neural tracking scores averaged across electrodes in the ROI (dots) and the mean of each age group (diamond). Boxplots show the median (horizontal line), 25th and 75th percentiles (box edges), and ±1.5 interquartile range (whiskers). No significant age-group difference in the neural tracking of acoustic envelope and onsets was found. D) Beta weights of TRFs averaged across frequency bands, ROI electrodes, and participants in each group for each acoustic predictor. Ribbons represent ±1 standard error. TRF time courses did not differ significantly between the two groups.

Figure 4B depicts the average neural tracking across the scalp for each age group. Mass-univariate independent-samples *t*-tests suggested that there were no clusters of electrodes with significant age-group difference in the neural tracking for both the envelope (*t_max_* = 2.372, *p* = 0.616) and onset (*t_max_* = 2.523, *p* = 0.380) features. Even when considering only the electrodes in the region of interest (ROI; yellow dots in Fig. 4B), defined as those with neutral tracking significantly above zero, we still found no evidence that the neural tracking differed between the younger group (envelope: *M* = 1.067 × 10⁻^3^, *SD* = 1.612 × 10⁻^3^; onset: *M* = 6.741 × 10⁻^3^, *SD* = 4.219 × 10⁻^3^) and middle-aged group (envelope: *M* = 1.367 × 10⁻^3^, *SD* = 1.874 × 10⁻^3^; onset: *M* = 5.730 × 10⁻^3^, *SD* = 5.990 × 10⁻^3^), for both acoustic envelope (Welch’s *t*(37.791) = 0.562, *p* = 0.578, 95% CI[−7.794 × 10⁻^4^, 1.378 × 10⁻^3^]) and onsets (Welch’s *t*(33.257) = −0.635, *p* = 0.530, 95% CI[−4.250 × 10⁻^3^, 2.228 × 10⁻^3^]; Fig. 4C).

To further test if the two groups differed in the temporal dynamics of neural responses to the acoustic features, we extracted the average *β* weights of the TRFs in the ROI over time from the full model with both the onset and envelope predictors (Fig. 4D). The average TRF corresponding to the acoustic onsets demonstrated an abrupt peak between 0–100 ms followed by a shallow valley, while that of the envelope showed a less defined and more delayed peak. Similar TRF characteristics were observed in previous studies using these acoustic representations (22,23). Crucially, using generalized additive models, we found no significant difference between younger and middle-aged listeners in the TRF curves either. Together, these results provided no evidence that envelope-based acoustic features were represented better or worse in middle-aged adults’ cortical responses.

### Middle-aged and younger adults show differences in speech perception and peripheral auditory health but not cognitive performance

Speech processing requires a complex interaction between precise auditory encoding and cognitive resource utilization (41–43). To understand whether our age groups differed on either of these domains, we looked beyond the EEG data and inspected measures from a series of audiological and cognitive tests (Fig. 5). Even though all participants were considered to have clinically normal thresholds, middle-aged adults (*M* = 10.708, *SD* = 3.976) had significantly higher pure tone averages (PTA) across 1, 2, and 4 kHz than younger adults (*M* = 5.625, *SD* = 3.918; Welch’s *t*(40.361) = –4.251, *p* < 0.001, *d* = –1.288, 95% CI[–7.500, –2.667]; Fig. 5A). We also measured extended high frequency (EHF) thresholds at 12.5 kHz, which is not commonly collected in audiology clinics but is a marker for lifetime acoustic exposure that is associated with CND (44). Middle-aged adults exhibited elevated EHF thresholds at 12.5 kHz (*Mdn* = 15.00), which was statistically greater than the thresholds for younger adults (*Mdn* = –2.500, *z* = –4.559, *p* < 0.001). These findings suggested that our middle-aged adults showed some degree of subclinical hearing loss at thresholds less than 4 kHz and some evidence for putative CND.

**Figure 5.**
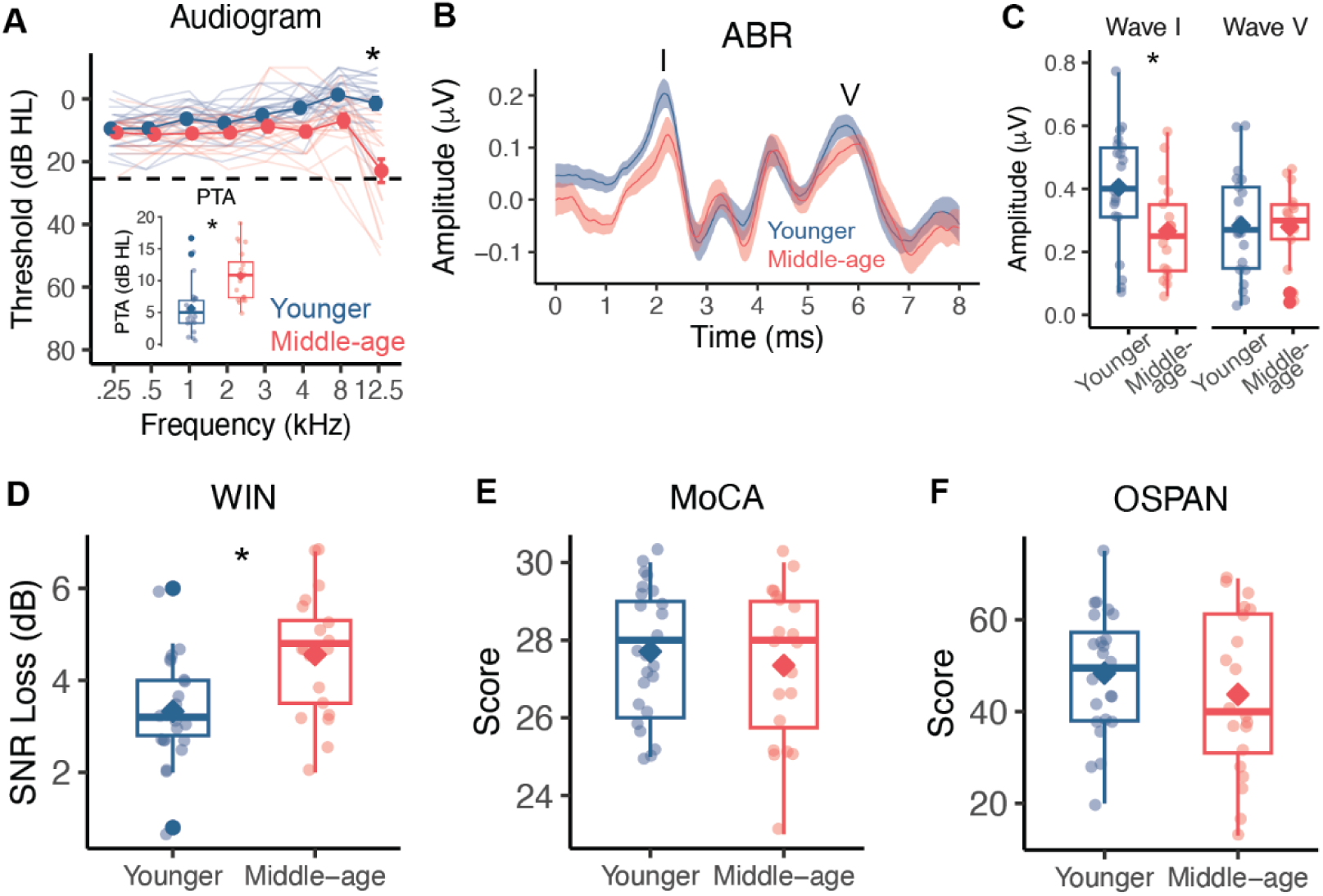
Results from audiological assessments, speech perception in noise performance, and cognitive tests. A) Hearing thresholds (dB HL) at frequencies from 0.25 to 12.5 kHz of individual participants (transparent lines) and group averages (opaque lines). Error bars reflect standard error of the mean. Pure tone averages (PTA) across 1, 2, and 4 kHz (inset) and thresholds at 12.5 kHz were significantly higher in middle-aged adults. B) Average auditory brainstem response (ABR) waveforms with the shaded region reflecting standard error of the mean. Wave I and wave V are marked by the I and V, respectively. C to F) Boxplots of individual ABR wave I and wave V amplitudes, Words in Noise (WIN) signal-to-noise ratio loss (SNR) in decibels (dB), Montreal Cognitive Assessment (MoCA) scores, and operation span task (OSPAN) scores for younger and middle-aged adults. Individual participants’ values (dots) and the group mean (diamond) are depicted. Boxplots show the median (horizontal line), 25th and 75th percentiles (box edges), and ±1.5 interquartile range (whiskers). Asterisks indicate statistical significance at alpha = 0.05 level.

Using the auditory brainstem response (ABR), we further probed the existence of CND in our middle-aged participants by examining ABR wave I and wave V amplitudes. Waves I and V of the ABR reflect synchronized activity in the auditory nerve (wave I) and central auditory structures, such as the lateral lemniscus (wave V) and inferior colliculus (wave V) (25,45). ABRs were originally proposed as a marker for CND in animal models, with smaller wave I amplitudes suggesting less integrity of the auditory nerve, and hence, CND (26). However, ABR wave I amplitudes can be highly variable between humans (46) and challenging to obtain (5). To combat these potential issues, we used gold-foil “tiptrodes” in the ear canal to improve the detection of ABR wave I and wave V amplitudes elicited to a 3 kHz tone. Middle-aged adults (*M* = 0.266, *SD* = 0.152) showed significantly lower wave I amplitudes compared to younger adults (*M* = 0.404, *SD* = 0.184; Welch’s *t*(35.979) = 2.536, *p* = 0.016, *d* = 0.819, 95% CI[0.028, 0.249]), providing evidence for putative CND in our middle-aged participants. In contrast, wave V amplitudes were not different between middle-aged (*M* = 0.279, *SD* = 0.130) and younger (*M* = 0.284, *SD* = 0.164) adults (Welch’s *t*(36.963) = 0.112, *p* = 0.912, *d* = 0.036, 95% CI[–0.090, 0.101]), suggesting that central auditory structures may increase neural responsiveness to compensate for degraded input from auditory nerve fibers.

One of the primary perceptual complaints of putative CND is difficulty understanding speech in noisy environments. To measure speech perception in noise performance, we administered the Words in Noise (WIN) test, which is commonly used in audiology clinics to measure word-level perception in multi-talker babble. The signal-to-noise ratio (SNR) loss, a metric of the SNR level required for accurate perception of words in background noise 50% of the time, was significantly higher in middle-aged adults (*M* = 4.505, *SD* = 1.339) compared to younger (*M* = 3.426, *SD* = 1.187) adults (Welch’s *t*(37.302) = –3.245, *p* = 0.002, *d* = –1.007, 95% CI[– 2.002, –0.463]). This indicated that the middle-aged adults in our study required quieter listening environments for accurate perception of words in noise than younger adults and further suggested that the diminished ABR wave I amplitudes may impact speech in noise.

While we observed several age group differences in audiological measurements, we did not observe differences in cognitive performance. All participants completed the Montreal Cognitive Assessment (MoCA) (47) as part of a prescreening measure and were required to score higher than 24 to participate in the study. Middle-aged (*M* = 27.350, *SD* = 1.954) and younger (*M* = 27.708, *SD* = 1.706) adults had comparable MoCA scores (Welch’s *t*(38.106) = 0.641, *p* = 0.525, *d* = 0.195, 95% CI[–0.773 1.489]), indicating normal cognitive function in both groups. Working memory, a cognitive process that is heavily involved in speech perception in noise, allows the listener to hold onto information, fill in missing phonemes and words, and continue to comprehend the speech stream (48). We administered the operation span task (OSPAN) (49) as a metric of working memory and executive functioning in our participants. As with the MoCA, we did not observe a significant difference in OSPAN scores between middle-aged (*M* = 43.750, *SD* = 17.586) and younger (*M* = 48.375, *SD* = 13.318) adults (Welch’s *t*(34.914) = 0.967, *p* = 0.340, *d* = 0.297, 95% CI[–5.081, 14.331]). Taken together, these tests provided evidence to indicate that our age groups were cognitively matched and that the observed differences in WIN and phoneme prediction accuracy from the PRP results may be driven at least partly by putative CND rather than cognitive processing.

### Putative CND and neural tracking of acoustics explain significant variance in phoneme prediction accuracy

Given the multitude of factors that can impact speech perception abilities, we probed the potential combined contributions of auditory and cognitive factors to cortical differentiation of phonemes. Specifically, we used a backward stepwise regression to calculate the variance in individual participants’ phoneme prediction accuracy from the PRP classification that could be explained by ABR wave 1 and wave V amplitudes, PTA, EHF thresholds at 12.5 kHz, neural tracking of acoustic envelope and onsets, OSPAN scores, and MoCA scores. Due to the normality assumption of the regression analysis, phoneme prediction accuracy values were first transformed into rationalized arcsine units (RAU) (50) and EHF thresholds were logarithmically transformed to normalize the distributions. Predictor variables were *z*-scored to allow for better interpretation of the standardized regression coefficients. The full regression model with all of the above listed predictors was not statistically significant (*F*_(8,28)_ = 2.289, *p* = 0.050), with an Akaike information criteria (AIC) of 109.73 and explained 22.3% of the variance in phoneme prediction accuracy. The results from the backward stepwise regression revealed that the combination of predictor variables that provided the lowest model AIC (AIC = 103.29) were ABR wave I amplitudes and neural tracking of acoustic envelope and acoustic onsets. This best fit model was statistically significant (*F*_(3,34)_ = 5.679, *p* = 0.003) with an adjusted-*R*^2^ of 0.275, explaining approximately 28% of the variance in phoneme prediction accuracy.

Interestingly, neural tracking of acoustic onsets (*β* = 1.830, *SE* = 0.808, *t* = 2.265, *p* = 0.030) and ABR wave I amplitudes (*β* = 2.461, *SE* = 0.655, *t* = 3.755, *p* = 0.001) were statistically significant predictors in the best fit model, suggesting that these two variables strongly contributed to phoneme prediction accuracy. Moreover, ABR wave I amplitudes had the largest coefficient, which not only indicated that ABR wave I amplitudes were the most important variable in explaining variance in phoneme prediction accuracy, but that smaller ABR wave I amplitudes were associated with lower phoneme prediction accuracy. This finding also provides evidence to suggest that participants with putative CND had noisier neural representations of phonemes, in line with the hypothesized contribution of putative CND to phoneme dedifferentiation. Neural tracking of the acoustic envelope (*β* = 1.583, *SE* = 0.784, *t* = 2.020, *p* = 0.051) was not a statistically significant predictor in the best fit model, but removing neural tracking of acoustic envelope resulted in a larger AIC (AIC = 106.13), indicating a poorer model fit without it. Together, these findings indicated that a combination of putative CND and neural tracking of acoustic information contributed to the neural representations of phonemes during continuous speech listening.

### Group differences in phoneme prediction accuracy remained after accounting for ABR wave I amplitudes and neural tracking of acoustic features

Given the significant proportion of variance in phoneme prediction accuracy explained by the predictors from the best fit regression model, we tested whether the age-group difference in phoneme prediction accuracy (in RAU) remained after controlling for these predictors. We addressed this with a residual analysis, in which we first ran a multiple linear regression model to predict phoneme accuracy using the independent variables that remained in the best fit model: ABR wave I amplitudes, neural tracking of acoustic envelope, and neural tracking of acoustic onsets. The residuals (i.e., differences from actual phoneme accuracy not explained by the predictors) were then extracted and compared across age groups. The results showed that phoneme prediction accuracy was still significantly lower for middle-aged adults compared with younger adults (*β* = −0.022, *SE* = 0.009, *t* = −2.478, *p* = 0.017). Therefore, although putative CND as suggested by smaller ABR wave I amplitudes significantly predicted phoneme encoding fidelity, this and the other predictors could not fully explain the age-group difference in phoneme prediction accuracy.

## Discussion

We recorded EEG responses to naturalistic speech from younger and middle-aged listeners with normal audiometric thresholds and assessed their cortical encoding of speech sound categories (Fig. 1). By training classifiers to predict phonemes from PRPs (i.e., average EEGs time-locked to phoneme instances), we found that phonemes were predicted significantly less accurately and with greater uncertainty for middle-aged adults compared with younger adults (Fig. 2). Information in the PRPs relevant to phoneme predictions was more broadly distributed across electrodes and showed a delayed peak timing in the middle-aged group (Fig. 2). Also, middle-aged adults’ PRPs aligned less with a phoneme representation based on the [syllabic], [sonorant,] and [continuant] features, suggesting a less specified featural relationship between phonemes (Fig. 3). Middle-aged adults also showed lower speech perception in noise performance (WIN score) and more pronounced signatures of putative CND (Fig. 5). Signatures of putative CND significantly predicted phoneme prediction accuracy from the PRP classifier and explained approximately one-third of the variance in the accuracy, jointly with neural tracking acoustic envelope and onsets. Notably, the age-group difference (middle-aged < younger) in phoneme prediction accuracy was still significant after controlling for peripheral and acoustic processing measures.

### Neural dedifferentiation to speech sounds in middle-age

Following (21) to derive PRPs, which represent cortical responses to individual phonemes, we obtained multiple lines of evidence in support of the hypothesized age-related neural dedifferentiation. The classifiers trained to predict phonemes from PRPs revealed less accurate and more uncertain predictions in middle-aged adults compared with younger adults (Figs. 2C and 2D). Furthermore, information relevant to phoneme predictions in the PRPs was delayed (Figs. 2F and 2I) and more distributed across electrodes (Figs. 2E and 2G), aligning with the less specialized and more correlated activity across the brain in older adults (17). Given that phoneme representations emerge from STG neural ensembles tuned to specific phonetic features, as revealed by ECoG data (9), we also explored the featural relationship between phonemes in our non-invasive PRP measures. We considered three features: [syllabic], [sonorant], and [continuant], which capture key manner class distinctions important to speech (e.g., vowels versus consonants, which are [+syllabic] and [-syllabic], respectively) and modeled the relationship between phonemes at an abstract level. The representational similarity analysis revealed that such a relationship was less distinctly represented in middle-aged adults’ PRPs when including all the three features (Fig. 3). Note that as the stimuli were presented in quiet, the results cannot be attributed to differential effects of noise masking on the two age groups. Together, these findings suggest that cortical representations of phonemes and the features that comprise them are less differentiated, or “fuzzier,” in middle-aged listeners. Thus, the current study not only replicated the neural dedifferentiation (8) in the context of meaningful, naturally produced speech sound units, but also offered novel evidence that such deficits may emerge as early as in middle-age.

The fuzzier, delayed, and more distributed network of phoneme representations may have significant downstream speech perceptual consequences. Age-related neural dedifferentiation has been linked to slower speed with which individuals make judgments about the perceived stimuli (51,52). This coincides with the relatively delayed relevance peak of our middle-aged participants, suggesting that they may show slower behavioral discrimination of phonemes. Even when no explicit judgments about phonemes were required, the fuzzier and more dispersed phoneme representation network is likely to induce subtle challenges to word recognition and more effortful listening. Indeed, middle-aged adults demonstrated poorer words-in-noise perception, consistent with past work reporting similar speech perceptual decline for middle-aged adults, regardless of audibility (1). In response to the reduced phonemic distinctiveness, listeners may allocate more cognitive resources in discriminating phonemes during spoken language processing. Per the Ease of Language Understanding model (53), the additional resources required for listening may allow for less cognitive allocation for performing a concurrent task, or achieving a higher-level reasoning of the discourse. In fact, some accounts, such as the information degradation hypotheses (54), have attributed the chronic changes to resource allocation as having the potential to increase the risk of cognitive decline.

Middle-aged adults sometimes subjectively report speech perceptual difficulties while exhibiting clinically normal pure-tone audiograms, leading to the perception that they are “overestimating” their difficulties (1,55–57). However, our findings challenge this view, suggesting that even without measurable hearing loss, middle-aged adults still struggle with maintaining distinct neural representations of critical, linguistically-relevant, information bearing units during spoken language processing. PRPs offer a promising non-invasive marker for detecting these subtle speech perception challenges. Although practical considerations and further development are necessary, incorporating PRPs into clinical protocols, alongside standard audiograms and other peripheral measures, could capture central decline in the representations of linguistically meaningful sound units.

Beyond clinical diagnostics, a key question raised by our results is the underlying mechanism(s) driving cortical dedifferentiation in middle-aged adults. Currently, we speculate that the dedifferentiation is driven by increased cortical noise due to changes in neurotransmitters. According to the computational model proposed by Li and colleagues (58,59), cognitive aging and neural dedifferentiation are linked to age-related changes to the availability of neurotransmitters. Specifically, decreases in neurotransmitter availability (e.g., dopamine) increase noise in the information processing of cortical neurons, which reduces the distinctiveness and fidelity of neural representations. Compared to younger adults, middle-aged and older adults are expected to demonstrate neural dedifferentiation given myriad evidence from positron emission tomography, revealing pervasive decline in markers of dopaminergic activity as a function of age across adulthood (60–62). In addition to dopamine availability, cortical noise may arise from decline in GABAergic inhibition. Studies in macaque models have noted dramatic age-related changes in the response patterns of the auditory cortex, such as increased spatial tuning and reduced temporal fidelity of response, which suggest a destabilized balance of neuronal excitation and inhibition (63). One driver of this imbalance could be lower levels of GABA, which reduces inhibitory control in the central auditory system and causes “central gain,” or maladaptive amplification of cortical responses (64,65). The hyperexcitability of neurons due to such gain has been observed in older adults compared with younger adults (66) and can lead to noisier, less distinct cortical representations that disrupt downstream speech perception (11,67).

Indeed, it is likely that the neural dedifferentiation of linguistically-relevant speech units— and the presumed cortical noise contributing to it—are inherited at least partly from subclinical peripheral deficits, particularly as a sequela of CND. Animal studies in mice offer direct evidence that CND increases internal noise in the auditory cortex, resulting in poorer behavioral detection of tones (12) and discrimination of speech sounds (68). In fact, central gain due to reduced GABA levels in older adults can arise out of the need to compensate for degraded afferent input due to auditory nerve atrophy (69). Thus, there may exist a relationship between CND and cortical dedifferentiation of phonemes, which is partially supported by our data.

### Neural dedifferentiation of phonemes is associated with CND, but peripheral and acoustic predictors only explain a third of the variance

We observed that more pronounced markers of CND were indeed associated with phoneme dedifferentiation. While CND has not been verified in humans *in vivo*, the wave I amplitude of the ABR is posited to reflect the integrity of the peripheral function, with reduced amplitude being a signature of putative CND (25,26,45). It is also argued that elevated thresholds at EHF (>8 kHz) mark underlying CND (70) and predict performance on speech perception in noise (71), although some (72) found no evidence for such predictive power after controlling for PTA at conventional speech frequencies. Our middle-aged adults did show signs of putative CND, as evidenced by reduced wave I amplitudes and higher EHF thresholds (Figs. 5A and 5C), despite normal audiograms. Notably, the stepwise regression analysis showed that lower ABR wave I amplitudes indicating putative CND were significantly related to lower phoneme prediction accuracy from the PRP classifier. With these findings and prior studies showing that central gain arises to compensate for peripheral deficits (69), we posit that similar compensatory mechanisms may be at play in middle-aged adults. That is, to adapt to degraded peripheral inputs induced by CND, disinhibition may occur throughout the central auditory system with the goal of increasing the neural gain. This is evidenced in our results as normalization of wave V amplitudes despite lower wave 1 amplitudes in middle-aged adults. Yet, while such adaptation reflects central plasticity, it may come at the cost of elevated cortical noise leading to neural dedifferentiation and poorer speech sound discriminability (68). Here, we further demonstrated that such costs include less differentiated phoneme representations.

In addition to markers of putative CND, the stepwise regression revealed that better cortical tracking of acoustic onsets was also associated with higher phoneme prediction accuracy. This association was compatible with the privileged role of onset information in cortical processing and speech recognition. Not only is the auditory cortex sensitive to acoustic onsets in speech (38,39), but the precise spatiotemporal patterns of cortical responses within milliseconds of a consonant onset robustly predict behavioral discrimination of the consonant (10). Computational models and behavioral data from the spoken word recognition literature also demonstrate an advantage for word-initial segments in guiding lexical access and managing lexical competition (73–75). Although acoustic onsets do not always coincide with word onsets, these findings highlight the importance of onset information in speech processing. Consequently, individuals whose brains are more capable of tracking acoustic onsets tend to retain more distinct phoneme representations.

Nevertheless, even after correcting for the effects of peripheral (wave I amplitudes) and acoustic (neural tracking of acoustic envelope and onsets) factors, the residual analysis showed that middle-aged adults’ lower phoneme prediction accuracy was still significantly different from young adults. The remaining accuracy gap between the two age groups raises the possibility that central factors of neural dedifferentiation exist which occur *independent* of peripheral changes and may relate proximally to speech perception. Future work integrating PRP, peripheral deficit measures, and concentrations of neurotransmitters in the auditory cortex could aid in uncovering these factors.

### Limitations

The mechanisms regarding neural dedifferentiation as discussed above are speculative, as we cannot verify that the PRPs derived from EEG relate directly to changes in the auditory cortex and especially the STG. However, the findings offer a few intriguing avenues for further research. These include examining age-related changes in phoneme or feature selectivity of neural responses in the STG using recording techniques with high spatial resolution. As mentioned, ECoG studies indicate that local sites in the STG tune to specific phonetic features (15,16). The degree of selective tuning can be quantified with measures such as the phoneme selectivity index (9) and the neural dedifferentiation hypothesis would then predict that such indices should decrease as a function of age. Furthermore, given the phonological theories of distinctive features (34) and the results from the representational similarity analysis (Fig. 3C), it may also be expected that age-related neural dedifferentiation would first affect finer featural distinctions lower in the phonological feature hierarchy, such as place distinctions, as compared with manner distinctions. Addressing these predictions could contribute to a more nuanced understanding of how aging relates to cortical changes in the central auditory system.

## Conclusion

In summary, we demonstrate that the neural distinctiveness of speech representations is reduced in middle-aged adults, who also demonstrate signatures of putative CND and speech perceptual deficits despite normal audiograms. While processing naturalistic speech, middle-aged adults show reduced speech sound distinction and featural relationships, delayed timing, more uncertainty, and more distributed processing relative to younger adults. These age-related changes cannot be entirely attributed to putative CND or lower-level acoustic processing. Fuzzier processing of the basic unit of spoken language processing may have significant downstream consequences on speech perception, such as elevated listening effort and greater need for cognitive-linguistic resources to scaffold spoken language processing. These findings highlight the need to consider age-related changes in cortical representations of critical speech sound categories and features when exploring the sources of speech perception challenges in middle-age.

## Materials and Methods

### Participants

Native English speakers were recruited from the Greater Pittsburgh area for a larger study on speech perception in noise abilities in middle-age. Here, we report data from the 44 participants who completed the continuous speech listening task from the larger speech perception in noise study. These included 24 younger-adults (22 females) aged 18–25 years (*M* = 21.417, *SD* = 2.020) and 20 middle-aged adults (12 females) aged 40–54 years (*M* = 46.050, *SD* = 4.347). All participants had: a) normal cognition, as determined by a score ≥ 25 on the Montreal Cognitive Assessment score (47); b) no severe tinnitus, as self-reported via the Tinnitus Handicap Inventory (76); and c) air conduction thresholds ≤ 25 dB hearing level (HL) at octave frequencies from 0.25 to 4 kHz. This study was approved by the University of Pittsburgh’s Institutional Review Board. All participants provided written consent and received monetary compensation for their participation.

### Audiological assessments

An otoscopic examination was first performed to ensure the ear canal and tympanic membrane were free of excess cerumen or other abnormalities. Air conduction thresholds were obtained using a MADSEN Astera audiometer, with Otometrics transducers (Natus Medical, Inc., Middleton, WI) and foam insert eartips in a sound attenuating booth. Thresholds were collected at 0.25, 0.5, 1, 2, 3, and 4 kHz. We also measured extended high frequency thresholds at 12.5 kHz using Sennheiser circumaural headphones and Sennheiser HDA 300 transducers. Most participants had binaural hearing thresholds ≤25 dB across standard test octave frequencies of from .25 to 8 kHz. One middle-aged participant had a single left ear threshold at 30 dB HL at 8 kHz, and one middle-aged participant had thresholds of 30- and 35-dB HL at 8 kHz. Binaural thresholds at 8 kHz for these two participants were 27.5- and 32.5-dB HL, respectively. For all other participants, binaural thresholds between 250–8000 Hz were ≤25 dB HL. Participants were not required to meet a particular threshold at the extended high frequencies to participate in this experiment.

Auditory brainstem responses (ABRs) were recorded using the Intelligent Hearing Systems (Miami, FL) Duet system. ABRs were collected to a 3 kHz tone burst while participants were seated in an electromagnetically shielded double-walled sound booth. Gold-foil tiptrode insert (reference/inverting) was placed in the right ear canal, and electrodes were placed at the forehead (ground) and Fz (active/non-inverting) location. Stimuli were presented at 86 dB nHL for 2048 sweeps to both the right and left ears, separately, using ER-3C transducers (Etymotic Research, Elk Grove Village, Illinois). The stimuli were sampled at 25 µs at a rate of 9.3 per second with alternating polarity. ABR wave I and wave V peak-to-trough amplitudes were manually marked by a trained audiology student on our research team and confirmed by a second expert member. Any inter-scorer disagreements between the two raters were settled through reviewing together.

### Speech perception in noise testing

We tested word-level speech perception in noise abilities using the Words in Noise (WIN) task (29). The WIN stimuli were comprised of spectrotemporally short with a shorter time window of integration, consisting of open-set single words with no surrounding linguistic context. The WIN test list consisted of seven groupings of five words masked in four-talker babble at the following signal-to-noise ratio (SNR) levels: 24, 20, 16, 12, 8, 4, and 0 dB. The words were presented in descending SNR level, resulting in more difficulty as the task progressed. Participants were instructed to repeat the word back to the best of their ability and to make their best guess if they were unsure. Two test lists were presented, resulting in 10 target words per SNR level. The SNR loss in dB was calculated for each subject, which reflects the optimal SNR level necessary to achieve 50% accuracy, using the equation: 26 – (*n* × 0.4), where *n* reflects the sum of words correctly identified (77).

### Continuous speech stimuli

The stimuli were continuous speech from the public domain audiobook *Alice’s Adventures in Wonderland* (78), recorded at 22.05 kHz sampling rate by a male American English speaker. Long pauses were reduced to a maximum of 500 ms and the audiobook was partitioned into segments of ∼60 s (range: 59–65 s). Participants listened to 15 segments in each of three listening conditions (quiet, speech-shaped noise, and reversed talker babble), in counterbalanced orders with the 15 segments within each condition presented chronologically to preserve the storyline. At the end of each segment, participants answered two four-choice comprehension questions to assess their understanding of the content in the preceding segment. The Montreal Forced Aligner (79) was run to obtain word and phoneme segmentations of the stimuli. In the current study, we analyzed and reported the data from the quiet condition.

### EEG acquisition and preprocessing

Continuous speech stimuli were presented binaurally through ER-3C insert earphones (Etymotic Research, Elk Grove Village, Illinois) at approximately 80 dB SPL. Electrophysiological responses to continuous speech were amplified and digitized with BrainVision actiCHAMP amplifier and collected using BrainVision PyCorder 1.0.7 (Brain Products, Gilching, Germany) with 64-channel actiCAP active electrodes (Brain Products) secured in an elastic cap (EasyCap; http://www.easycap.de/). Electrodes were placed on the scalp according to the International 10-20 system (80) and a common ground was placed at the AFz electrode site. Electrode impedance was less than 25 kΩ for all channels. Raw EEG responses were recorded at a 25 kHz sampling rate and preprocessed using EEGLAB 14.1.2 in MATLAB. The data were downsampled to 128 Hz for computational efficiency, bandpass filtered from 1 to 15 Hz using minimum-phase causal windowed sinc FIR filters, and re-referenced to the average of two mastoid electrodes (81–83). At the re-referenced channels, electrical activity outside the ±3 standard deviation range of the surrounding channels was rejected and interpolated, and artifacts were suppressed with artifact subspace reconstruction (ASR) (84). The ASR-cleaned data were epoched from −5 to 70 s relative to the story segment onset. Independent component analysis was performed on the epoched data to remove ocular and muscular artifacts and reconstruct the EEGs.

### Phoneme-related potential analyses

We adopted the phoneme-related potential (PRP) approach (21) to examine cortical representations of phoneme categories. For each phoneme, processed EEGs time-locked to 0.0–0.5 s after the onset of each phoneme instance were averaged to create a 61 (electrodes) × 64 (time points) PRP array representing the typical evoked response to that phoneme. Due to highly skewed phoneme frequency distribution in natural speech (e.g., the schwa AH accounted for nearly 10% of all instances while OY accounted for only 0.08% in our stimuli), we retained only the 31 phonemes with >319 instances (i.e., >1% of instances of all phonemes) excluding the highly frequent AH. To further minimize the possibility that phonemes with many instances could lead to high signal-to-noise ratios in the PRPs and bias the results, we imposed an upper limit on the number of instances included in the PRP calculation, which was set to 319. For phonemes with over 319 instances, we averaged a random subset. Subsequent analyses showed that PRP decoding accuracy was not significantly correlated with phoneme frequency in the audiobook (*r* = −0.139, *p* = 0.456).

#### PRP classification

We used EEGNet (31) to predict phoneme classes from PRPs. EEGNet is a compact convolutional neural network that has been demonstrated to learn useful features for classifying single-trial EEGs and event-related potentials across different brain-computer interface paradigms while allowing interpretation of the learned features. Here, we used the EEGNet-8,2 model with eight temporal filters, two spatial filters per temporal filter, a dropout rate of 0.5 (31). We trained the model on the PRP datasets from younger and middle-aged adults separately with 20-fold cross-validation. At each iteration, 5% of the data were held out as the test set and a random 15% of the non-test data served as the validation set. The model was trained on the remaining PRPs with an initial learning rate of 0.01 and a batch size of 16 for a maximum of 300 epochs to minimize cross-entropy loss. The learning rate was decreased by a factor of 0.7 every 100 epochs. The model at the epoch with the lowest validation loss was selected to predict the test set. A prediction was correct when the highest-probability phoneme matched the actual phoneme, and the prediction uncertainty was quantified as the Shannon entropy of the probabilities over all phonemes: −∑ *p*⋅ln(*p*) (higher indicated more uncertain).

The whole training and testing, as well as the selection of phoneme instances in PRP derivation, was repeated 20 times with different random seeds to ensure robust findings. Additionally, the younger group had four more participants than the middle-aged group, we randomly dropped four younger adults at each repetition to prevent any biases due to the slightly imbalanced data. The accuracy and uncertainty results were averaged across all repetitions and phonemes for each participant and compared across the two age groups (see Statistical analyses for details of all statistical tests). Model training was conducted using *PyTorch* 2.1.0 and the EEGNet used the implementation in *TorchEEG* (github.com/torcheeg).

#### Feature relevance

Despite neural network’s black-box reputation, EEGNet allows interpretations of its learned representations. We applied the Deep Learning Important FeaTures (DeepLIFT) algorithm (32) with the Rescale rule implemented in the Pytorch library *Captum* (85) to assess which features (here, responses at a particular electrode or time point in a PRP) were relevant for the trained EEGNet models to correctly predict the target phonemes of the test-set PRPs. DeepLIFT assigned a score representing the amount of evidence each input feature provided toward the target phoneme by backpropagating the contributions through the network to the input. The algorithm yielded a 61 × 64 contribution score matrix for each PRP, with positive and negative values denoting evidence for and against the target phoneme, respectively, and zero indicating irrelevance. We focused on quantifying relevance regardless of the sign of evidence by taking the absolute scores and *z*-transformed the scores within each matrix to compute relative relevance. The results were averaged across phonemes and across the 20 repetitions to obtain, for each participant, a single score matrix summarizing the relevance of each electrode over time. We averaged across time and computed the variance between electrodes as a measure of relevance dispersion (smaller indicates more distributed relevance). We also averaged across electrodes to determine the timing of the relevance peak relative to PRP onset. A mass-univariate independent *t*-test was run to compare the relevance topographies of the two age groups and identify clusters of electrodes showing a significant group difference.

#### Representational similarity analysis

The representational similarity analysis (35) compared the PRPs from the PRP classifier training (averaged over all repetitions) against a featural description of the phoneme relationship according to phonological theories (34). Three binary distinctive features were considered: [syllabic], indicating whether a sound can be the syllable nucleus; [sonorant], indicating whether a sound is produced with an open vocal tract and continuous non-turbulent airflow; [continuant], indicating whether a sound is produced with oral cavity blocking. The presence and absence of each feature were coded as 1 and 0, respectively. Based on the feature representation, we used the Python *rsatoolbox* library (rsatoolbox.readthedocs.io) to construct a representational dissimilarity matrix (RDM) containing pairwise phoneme distances with the Euclidean distance as the dissimilarity measure. The same procedure was followed to obtain RDMs for reduced models including [syllabic] and [sonorant] or only [syllabic].

The analysis was similarly applied to PRPs to derive neural RDMs capturing the neural dissimilarities between phonemes over time, which were each compared with the feature RDM of interest through Spearman’s correlation *ρ*. Higher *ρ* suggests better featural-neural representational alignment. As the electrodes might not all bear critical feature information, we employed a forward selection algorithm to determine the optimal set of electrodes for each age group. First, we ranked the electrodes based on the relevance scores from the relevance analysis. Next, the top two most relevant electrodes were included to build neural RDMs and calculate a *ρ* value averaged over time and over participants in the group, which represented the overall fit to the feature RDM. This process iterated, each time adding the most relevant electrode from the rest of the set, until the overall *ρ* stopped improving. The optimal *ρ* curves of the younger and middle-aged listeners were compared using generalized additive modelling. RDMs were also constructed from the prediction confusion matrices of the PRP classifiers and compared with the full feature RDM using the same *ρ* metric. The observed *ρ* difference between the younger and middle-aged groups was tested using permutation analysis.

### Estimation of neural tracking of acoustics

The continuous spectro-temporal acoustic properties of the stimuli were modelled as spectrograms using gammatone filters, which stimulate the frequency analysis performed in the cochlear (86,87). We used the gammatone filters from the Python *Eelbrain* package (88) with 256 filter channels and cut-off frequencies of 0.02–5 kHz. The 256-band spectrograms were downsampled to 1 kHz, scaled with an exponent of 0.6, and summed into eight logarithmically spaced frequency bands to derive 8-band gammatone spectrogram predictors for subsequent analyses. Additionally, given that responses in the auditory cortex are sensitive to acoustic onsets (38,39), we controlled for representations of acoustic onsets by applying an auditory edge detection model (89) to the 256-band gammatone spectrograms using the parameters in Brodbeck et al. (86). The resulting onset spectrograms were summed into eight logarithmically spaced frequency bands, creating 8-band onset spectrogram predictors.

Cortical tracking of acoustics was estimated by fitting multivariate temporal response function (mTRF) models (40) that describe the linear forward mapping from the acoustic predictors to EEGs. TRFs to a continuous stimulus such as the gammatone spectrogram can be interpreted as the evoked responses to an elementary event at each frequency band of the spectrogram. We included either or both the gammatone spectrogram and onset spectrogram as predictors and used the boosting function in *Eelbrain* to derive TRFs in each electrode at time lags from –100 to 500 ms with L1-error following a 5-fold cross-validation procedure. At each iteration, EEGs from three of the 15 story segments were held out as the test data while those from the other segments were used to train and validate TRF models, which were averaged to predict the test set when the estimation was finalized. The Pearson’s correlation coefficient (*r*) between the predicted and observed EEGs measured how accurately the brain tracked the predictor. We estimated neural tracking that could be uniquely attributed to the predictor of interest (Δ*r*) by taking the *r* difference between the full model including both spectrogram and onset spectrograms and the model with that predictor left out. Topographies of neural tracking were compared across the two age groups with a mass-univariate independent *t*-test. They were also compared against zero with a mass-univariate one-sample *t*-test to identify the region of interest (ROI), or electrodes with a significantly greater-than-zero Δ*r.* A single neural tracking score was obtained for each participant by average the Δ*r* values in the ROI.

### Statistical analyses

Welch’s two sample *t*-tests were calculated using the *rstatix* package (90) in R (version 4.3.1) (91) to compare the two age groups across variables that were normally distributed. Normally distributed variables included audiological and cognitive tests, relevance dispersion, and average neutral tracking (Δ*r*) of acoustic envelope and onsets in the ROI. Normality was confirmed with Shapiro-Wilks tests (*p* > 0.05) and visual inspection of quantile-quantile plots. Mann-Whitney *U* tests were used to compare phoneme prediction accuracy and uncertainty values from the PRP classifier, latencies of relevance peak, and EHF thresholds at 12.5 kHz between groups because the data were not normally distributed.

A backward stepwise regression was performed to determine which factors explained the most variability in phoneme prediction accuracy. Because regressions have no non-parametric alternatives, variables which violated the normality assumption were transformed. Phoneme prediction accuracy was converted to rationalized arcsine units (RAU) (50) to normalize the distribution and mitigate floor and ceiling effects. EHF thresholds at 12.5 kHz were logarithmically transformed to also meet normality after adding 25 to all thresholds to ensure no negative or non-zero values. Then, a multiple regression was fit using the *lm* function in *lmerTest* (92) with RAU phoneme prediction accuracy as the outcome variable and predictor variables of ABR wave I amplitudes, ABR wave 5 amplitudes, PTA across 1, 2, and 4 kHz, transformed EHF thresholds at 12.5 kHz, neural tracking of acoustic onsets, neural tracking of acoustic envelope, OSPAN scores, and MoCA scores. The backward stepwise regression was performed using the *stepAIC* function in the *MASS* package (93) in R. The backward stepwise regression removed a predictor variable at each step to find the combination of variables that resulted in a model with the lowest Akaike Information Criteria (AIC). In the residual analysis, the predictors in the best fit model according to the stepwise regression were included in a multiple regression model using the *lm* function to predict RAU phoneme prediction accuracy. The *residuals* function was used to extract the residuals, which met the normality assumption and were compared across age groups with a Welch’s *t*-test.

For topographical data, mass-univariate *t*-tests implemented in *Eelbrain* were conducted to compare variables of interest against zero (one-sample tests, one-tailed) or between younger and middle-aged adults (independent-samples tests, two-tailed). The mass-univariate *t*-test is a cluster-based permutation test that uses a *t*-value equivalent to uncorrected *p* ≤ 0.05 as the cluster forming threshold. Clusters were based on the identification of meaningful effects across groups of adjacent electrodes that showed the same effect (33). A corrected *p*-value was then computed for each cluster based on the cluster-mass statistician null distribution from 10,000 permutations (33). The largest absolute *t*-value from the cluster (*t*_max_) is reported as an estimate of effect size (94). The tests were used to compare the topographical scalp maps of neural tracking of the acoustic predictors and the time-varying relevance scores (analysis time window: 0 to 500 ms). A mass-univariate *t*-test was also run on the data from all participants to identify the ROI for acoustics tracking, which was the set of electrodes with tracking scores significantly greater than zero.

Generalized additive models using the *mgcv* package (95) in R were performed to identify group differences in the series data, which included the *ρ* values from the RDM analyses and the average beta weights in the ROI from the acoustic tracking mTRFs. The models consisted of the constant difference between the two groups and the smooth term s(time, by = group) that captured potential nonlinearity over time while allowing the degree of nonlinearity to vary across groups. We also included a smooth term, s(time, subject, bs = “fs”, m = 1), to allow differing degrees of nonlinearity across participants. We plotted the estimated group difference curve and identified the interval during which the difference was significant (when 95% confidence interval excluded zero).

The permutation test for the observed group difference in *ρ* from the RDM analyses was conducted by randomly shuffling the rows of the confusion matrices, computing a new *ρ* difference, and repeating the process for 5,000 times to build a null distribution of differences. The proportion of difference values smaller than the observed one was treated as the *p*-value.

## Acknowledgements

This research used the resources provided by the Quest high-performance computing cluster at Northwestern University and the University of Pittsburgh Center for Research Computing, RRID:SCR_022735 (NSF OAC-2117681). The authors would like to thank Sarah Anthony, Olivia Flemm, Claire Mitchell, and Leslie Zhen for their assistance with participant recruitment and data collection.

## Funding

This research was funded by National Institutes of Health Grant F31DC020085 to JRM, National Institutes of Health grant R21DC018882 to AP, National Institutes of Health grant R01DC013315 to BC, and the PNC-Trees Charitable Trust to BC and AP.

## Competing interests

The authors declare they have no competing interests that are relevant to the content of this article.

## Author Contributions

Conceptualization: Zhe-chen Guo, Jacie R. McHaney, Aravindakshan Parthasarathy, Bharath Chandrasekaran

Data Curation: Zhe-chen Guo, Jacie R. McHaney Formal Analysis: Zhe-chen Guo, Jacie R. McHaney

Funding Acquisition: Jacie R. McHaney, Aravindakshan Parthasarathy, Bharath Chandrasekaran

Investigation: Jacie R. McHaney

Methodology: Zhe-chen Guo, Jacie R. McHaney, Aravindakshan Parthasarathy, Bharath Chandrasekaran

Project Administration: Jacie R. McHaney, Aravindakshan Parthasarathy, Bharath Chandrasekaran

Resources: Aravindakshan Parthasarathy, Bharath Chandrasekaran

Software: Zhe-chen Guo, Jacie R. McHaney

Supervision: Bharath Chandrasekaran

Validation: Zhe-chen Guo, Jacie R. McHaney, Bharath Chandrasekaran

Visualization: Zhe-chen Guo, Jacie R. McHaney

Writing – Original Draft Preparation: Zhe-chen Guo, Jacie R. McHaney, Bharath Chandrasekaran

Writing – Review & Editing: Zhe-chen Guo, Jacie R. McHaney, Aravindakshan Parthasarathy, Bharath Chandrasekaran

## Notes

### Competing Interest Statement

The authors have declared no competing interest.

